# Rapid motility arrest of the Lyme disease spirochete, *Borrelia burgdorferi*, by vaccine-induced OspA antibodies

**DOI:** 10.64898/2026.07.24.740573

**Authors:** Ankita Bhattacharyya, Raymond Yau, Thomas M. Bartlett, Nicholas J. Mantis

**Author notes:** **Corresponding author:** Dr. Nicholas J Mantis, 120 New Scotland Ave, Albany, NY 12208; **email:**.

## Abstract

Outer surface protein A (OspA)-based Lyme disease vaccines elicit antibodies that inhibit transmission of infectious *Borrelia burgdorferi* spirochetes from ticks to humans, although the mechanism by which this occurs is unclear. Here we demonstrate using high-resolution, single-cell fluorescence video microscopy that *B. burgdorferi* motility is arrested within minutes by human OspA-specific monoclonal antibodies and polyclonal OspA antisera from mice immunized with a candidate OspA mRNA lipid nanoparticle vaccine.

---

*Borrelia burgdorferi*, the spirochetal agent of Lyme disease, is transmitted to humans via the black legged tick (*Ixodes scapularis*). Lyme disease vaccines based on <u>O</u>uter <u>S</u>urface <u>P</u>rotein <u>A</u> (OspA) elicit antibodies that when ingested by *I. scapularis* during a blood meal entrap *B. burgdorferi* within the tick midgut, thereby inhibiting transmission ^1, 2, 3^. OspA-specific mouse (e.g., LA-2) and human (e.g., 221-7) monoclonal antibodies (mAb), when passively administered to mice or non-human primates prior to tick challenge also interfere with *B. burgdorferi* transmission, even in the absence of complement and Fc receptor (FcR) activities ^4, 5, 6, 7^. Some of those same mAbs were recently shown to inhibit *B. burgdorferi* transmigration across a 3-micron porous membrane in a Transwell unit, leading us to postulate that OspA antibodies alter spirochete motility ^8^. However, conclusions regarding the direct effects of OspA antibodies on *B. burgdorferi* motility at the single cell level were confounded in the Transwell assays by the fact that OspA antisera and certain mAbs induce *B. burgdorferi* agglutination and large multicellular aggregates with the potential to impede bacterial passage through the Transwell (3 micron) membrane ^8, 9^.

To resolve this issue, we developed a hanging drop assay that enabled us to monitor the motility of *B. burgdorferi* at the single-cell level. Specifically, one microliter of mid-log phase spirochetes in BSK-II medium containing 2% gelatin was spotted onto a glass coverslip, which was then inverted and placed on a depression slide. The slide was mounted on an inverted microscope for timelapse imaging. Individual cells in each field of view were masked and tracked using the TrackMate plugin in Fiji to determine their mean track speed (µm/s), and distance traveled (µm) over the corresponding tracking duration (≤2 min) (see Materials and Methods). An OspA^+^ *B. burgdorferi* HB19-R1 strain expressing mScarlet (GGW1073) and an OspA^-^ *B. burgdorferi* strain expressing GFP (GGW1082) were mixed 1:1 for the assay and were visualized in DSRed (Ex 554/23 nm, Em 609/54 nm) and GFP (Ex 466/40 nm, Em 525/50 nm) channels. The strains traveled at an average speed of 1.4 ± 0.1 µm/s (**Figure 1A; Video 1**). This value is slightly lower than those reported for *B. burgdorferi* strain 297 (4.2 µm/s) but essentially identical to that reported by others using strain JD1 ^10, 11^. The speeds and distances traveled were consistent across the different timepoints evaluated (<5, 30, and 60 min). At the cell concentrations tested, there was no measurable degree of spirochete auto-aggregation or visible clumping, thereby facilitating single cell tracking efforts. A wide variety of dynamic spirochete morphologies likely associated with translocation and flexing as described by others were noted in the videos (**Table S1; Videos 2-6**) ^10, 12, 13, 14^.

**Figure 1.**
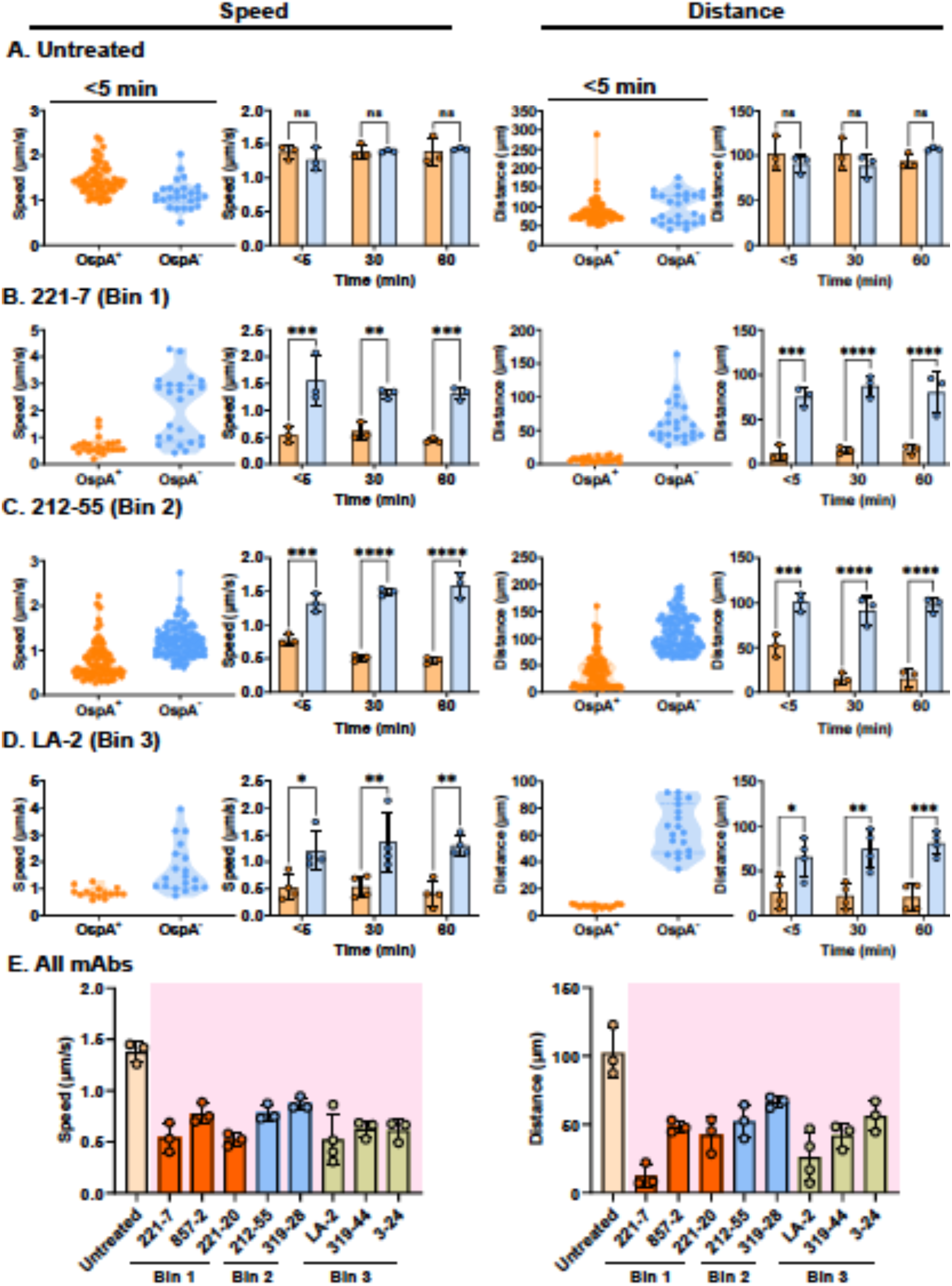
*B. burgdorferi* motility arrest in response to OspA mAbs, as indicated by reduced velocity and supported by reduced distance traveled. GGW1073 (OspA^+^; mScarlet) and GGW1082 (OspA^-^; GFP) spirochetes were mixed in a 1:1 ratio and treated with (**A**) PBS, (**B**) 221-7, (**C**) 212-55, or (**D**) LA-2 then imaged as described in Materials and Methods. Cells were masked and tracked using TrackMate in Fiji to measure their speed (µm/s) and total distance traveled (µm). Violin plots display speed/distance traveled of OspA^+^ spirochetes compared to the OspA^-^ controls at the <5 min timepoint for a single experiment in which each datapoint indicates the motility of a single spirochete. The bar graphs depict the average speed and distance of OspA^+^ and OspA^-^ spirochetes at <5 min, 30 min, and 60 min of mAb treatment for three biological replicates. Statistical significance was determined by Two-way ANOVA followed by Šídák’s multiple comparisons test. Asterisks indicate degrees of significance (*, *P*≤0.05, **, *P*≤0.01; ***, *P*≤0.001; ****, *P*≤0.0001; ns, not significant). (**D**) Speed and distance traveled of OspA^+^ spirochetes treated with each of the eight mAbs tested. Statistical significance was determined by One-way ANOVA followed by Dunnett’s multiple comparisons test. Pink shading indicating *P*≤0.05 as compared to untreated spirochetes.

To determine the compatibility of the hanging drop assay to measure antibody-mediated changes in spirochete motility, we performed a pilot study in which a 1:1 mixture of fluorescently tagged *B. burgdorferi* OspA^+^ and OspA^-^ strains were treated with the transmission blocking mouse mAb, LA-2 IgG, then tracked using single-cell microscopy. We observed that LA-2-treated *B. burgdorferi* OspA^+^ spirochetes stopped swimming within a matter of minutes and that changes in both speed and distance traveled were measurable under these conditions (**Video 7**). Thus, the hanging drop assay affords a means of quantitating *B. burgdorferi* distance traveled and speed following antibody treatment.

We therefore investigated *B. burgdorferi* motility following exposure to a panel of human mAbs targeting conserved (bin 1) and variable (bins 2-3) B cell epitopes on OspA (**Table 1**) ^15, 16^. Among these, mAb 221-7 (bin 1) was of particular interest, as it has been shown to protect both mice and non-human primates against tick-transmitted *B. burgdorferi* infections ^5, 6, 16^. It was also one of the most effective mAbs at inhibiting *B. burgdorferi* transmigration *in vitro* ^8^.

**Table 1.**
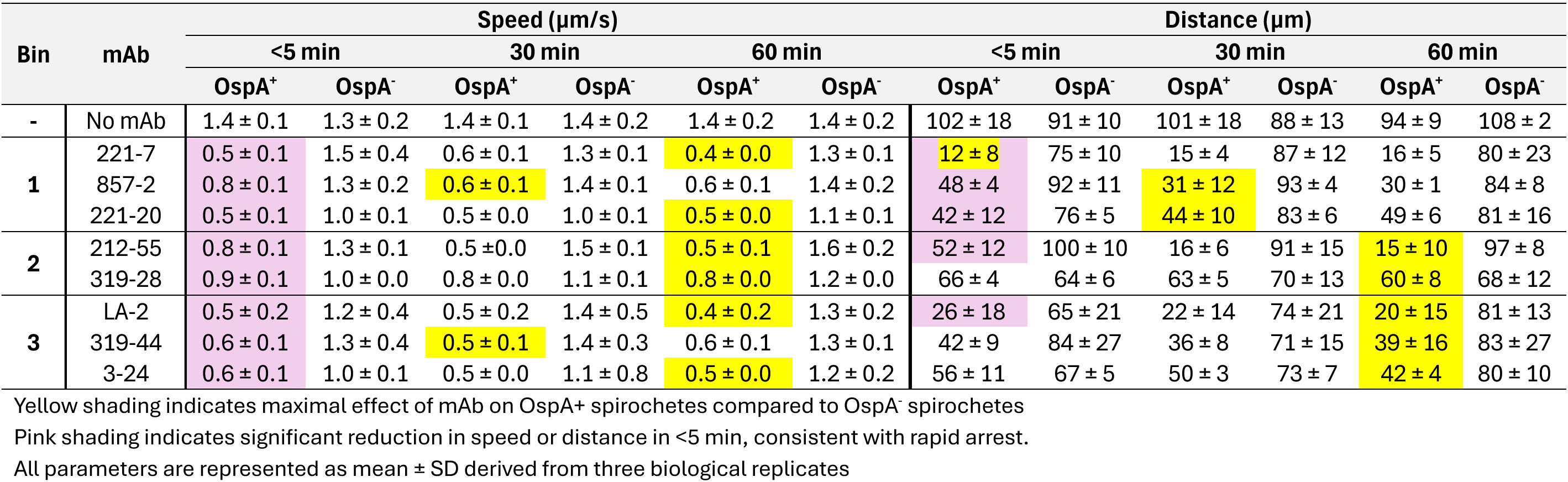
Motility parameters (speed and distance traveled) of OspA^+^ (GGW1073) and OspA^-^ (GGW1082) spirochetes in untreated and mAb-treated conditions after <5, 30, and 60 min of incubation.

In the hanging drop assay, 221-7 (10 µg/mL) reduced the speed of strain GGW1073 (OspA^+^) >3-fold relative to GGW1082 (OspA^-^) (0.5 ± 0.1 µm/s versus 1.5 ± 0.4 µm/s) within 5 min (**Figure 1B; Video 8**). Correspondingly, the total distance covered by OspA^+^ spirochetes in the presence of 221-7 was only 12 ± 8 µm compared to 75 ± 10 µm for OspA^-^ spirochetes. There was no further reduction in speed or distance traveled at 30 or 60 min timepoints, demonstrating that 221-7’s effects on *B. burgdorferi* motility were immediate and lasting. Two other Bin 1 mAbs, 857-2 and 221-20 whose epitopes overlap with 221-7, also inhibited *B. burgdorferi* B31 motility within a matter of minutes but to a lesser degree than 221-7 (**Table 1; Figure S1; Videos 9, 10**).

The remaining four human OspA-specific mAbs in our collection targeting epitope bins 2 (212-55, 319-28) and 3 (319-44, 3-24) also impacted *B. burgdorferi* B31 motility, albeit to varying degrees and at different time points (**Table 1**; **Figure 1; S1; Videos 11-15**). For example, while 212-55, 319-44 and 3-24 reduced *B. burgdorferi*’s motility in <5 min, maximal impact did not occur until 60 min. LA-2 was among the more potent mAbs, as it reduced the speed and distance covered by OspA^+^ spirochetes to 0.5 ± 0.2 µm/s and 26 ± 18 µm, respectively, within <5 min (**Figure 1D, Video 13**). These results demonstrate that both mouse and humans mAbs targeting a variety of epitopes on OspA have immediate and lasting effects on spirochete motility *in vitro*, possibly accounting for their ability to inhibit transmission of *B. burgdorferi in vivo*.

To determine if vaccine-induced OspA immune sera also interferes with spirochete motility, serum samples from groups of mice that had received an OspA mRNA vaccine across a range of doses (1-0.01 µg) were applied to GGW1073 (OspA^+^) spirochetes and motility was monitored by microscopy ^17^. Spirochete motility was unchanged when GGW1073 was treated with sera from sham vaccinated mice (**Figure S2; Video 16**). On the other hand, sera from vaccinated mice reduced spirochete speed and total distance traveled in a manner that corresponded to vaccine dose (**Figures S2; Videos 17-20)**. Sera from the lowest vaccine dose group (0.01 µg) had little to no impact on spirochete motility while sera from high dose vaccinated animals affected speed (0.6 ± 0.1 µm/s) and distance (25 ± 4 µm). The impact of immune sera on spirochete motility *in vitro* correlated with observed protection against *B. burgdorferi* infection in the tick challenge model (**Figure 2, S3**). These data suggest that motility arrest may constitute a correlate of protection after vaccination for Lyme disease ^18, 19^.

**Figure 2.**
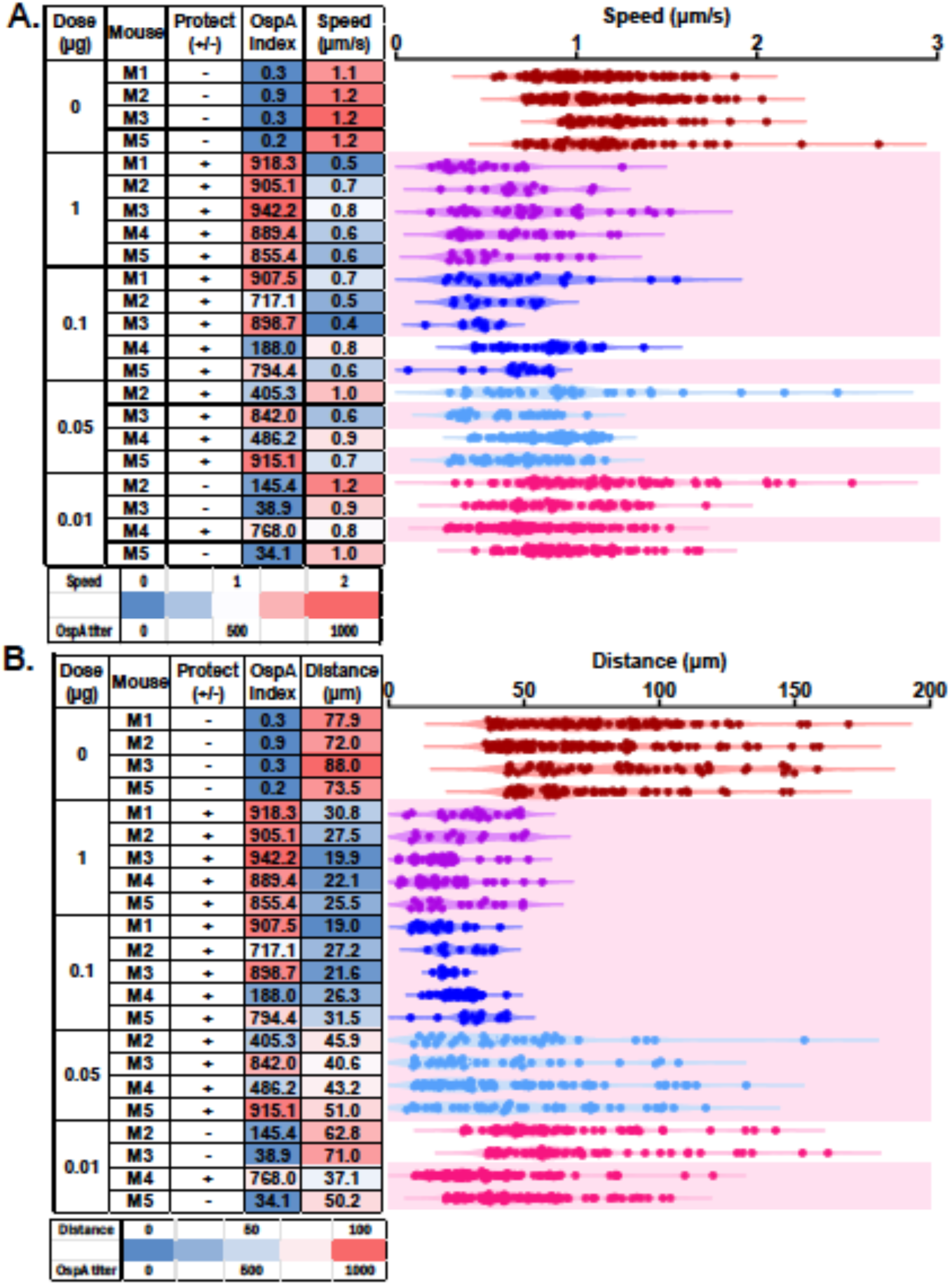
Impact of OspA antisera elicited by OspA mRNA vaccination on *B. burgdorferi* speed and distance traveled. GGW1073 (OspA^+^) spirochetes were treated with 1:50 dilution of antisera from mice that were sham vaccinated or vaccinated with a range of doses of an OspA serotype 1 mRNA-LNP vaccine, as described in Materials and Methods. Mouse protection from *B. burgdorferi-*tick challenge is reported as **+**: protected; **-**: unprotected (as determined previously by organ culture) {Finn, 2026 #8338}. OspA median fluorescence intensity (MFI) indices (as determined previously by serological analysis) are plotted as a heatmap, followed by heatmaps for the average (A) speed (B) distance traveled of all OspA^+^ spirochetes tracked, and violin plots where each datapoint indicates the respective motility parameter of each spirochete after 30 mins of treatment with the antisera from each vaccination group. Statistical significance was determined by One-way ANOVA followed by Dunnett’s multiple comparisons test. Pink shading indicates *P*≤0.05 compared to untreated spirochetes (not shown in figure).

While the mechanism by which OspA antibodies interrupt *B. burgdorferi* is not immediately clear, it is interesting to note that motility arrest induced by mAbs and immune sera was preceded by, or coincided with, the formation of alternate spirochete morphologies such as lariats, rings, and figure 8 that occurred in the absence of visible cell-cell agglutination (**Figure 3; Video 21**). Based on work by Barbour et al {Barbour, 1983 #7359}, we speculated that these morphologies might be due to antibodies driving OspA localization to the cells poles owing to mis-localization of cholesterol rafts and altering membrane fluidity. However, this was not borne out experimentally, as we did not observe OspA re-localization concurrent with spirochete looping, after tracking with LA-2 tagged with a fluorescence conjugate (**Video 22**). Hence, we concluded that although antibodies might drive OspA to spirochete poles, it is not necessary for the bacteria to loop on the ends and inhibit motility.

**Figure 3.**
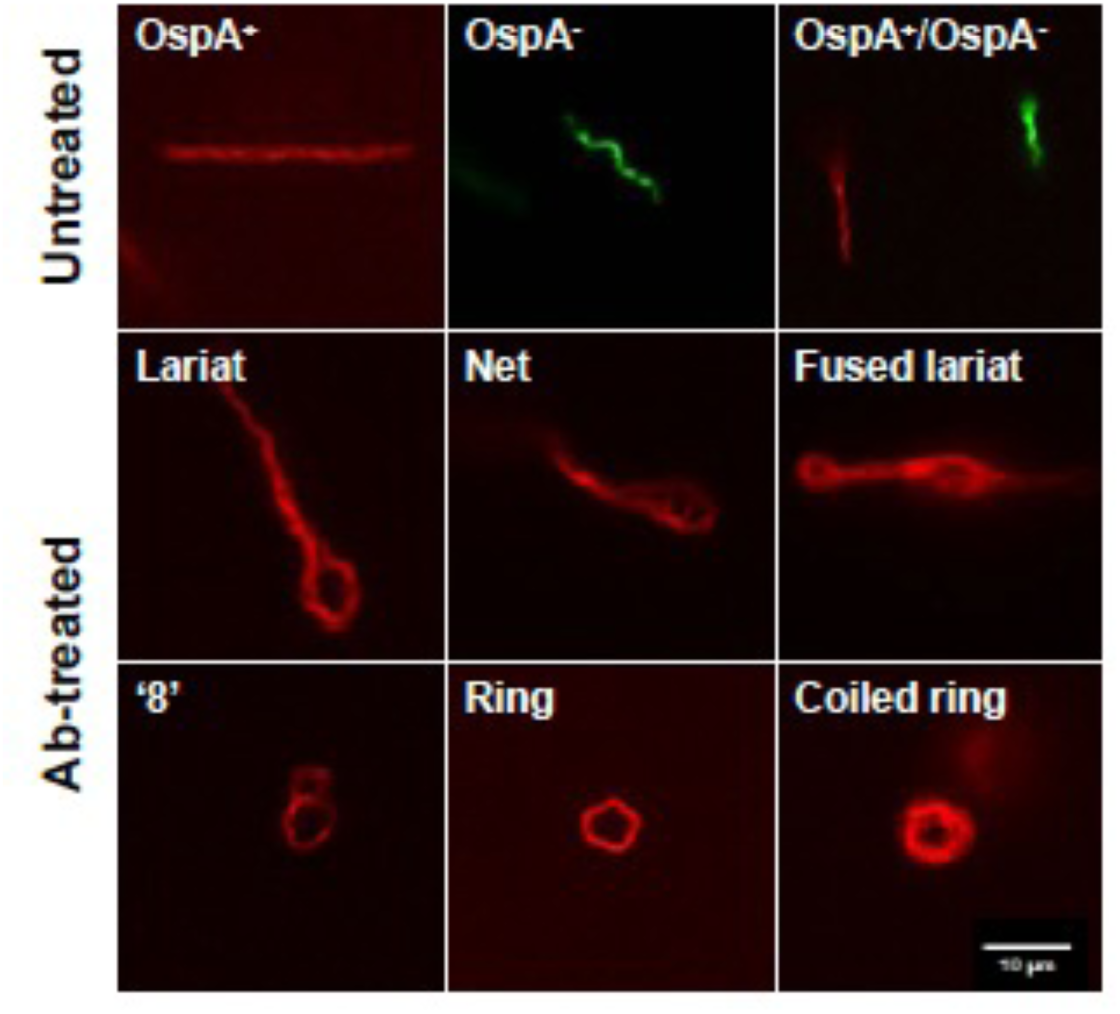
Single cell microscopy reveals altered spirochete structures when treated with OspA antibodies. GGW1073 (OspA^+^; mScarlet) and GGW1082 (OspA^-^; GFP) were mixed in a 1:1 ratio and were either left untreated or were treated with 10 µg/mL or 1:50 dilution of OspA mAbs or polyclonal sera respectively. While both strains showed their usual helical shape in untreated conditions, OspA^+^ strains altered morphologies in <5 min or 30 mins of treatment with OspA antibodies. Imaging was conducted using a Nikon ECLIPSE Ti2-E microscope equipped with a 40X fluorescence oil objective in the DSRed and FITC channels, respectively, as described in Materials and Methods.

In summary, we conclude that both OspA-specific mAbs and vaccine-induced immune sera rapidly and sustainedly reduce *B. burgdorferi* motility and translocation, as measured by mean track speed and supported by reduced distance traveled. This finding is significant in the context of how OspA antibodies may function within the tick midgut to limit spirochete penetration across the midgut epithelium and therefore transmission to vertebrate hosts. Moreover, motility arrest would also explain how OspA antibodies exert their transmission blocking activity in the absence of complement and FcR interactions ^4, 7, 8^.

## Materials and Methods

### *B. burgdorferi* strains and culture conditions

*B. burgdorferi* strain GGW1073 (OspA^+^) is a derivative of HB19-R1 transformed with an IPTG-inducible mScarlet-I reporter plasmid expressing OspA serotype 1 (ST1) under control of the P*_ospAB_* promoter ^16, 20^. GGW1082 (OspA^-^) is a derivative of HB19-R1 transformed with pTM61, which constitutively expresses a highly fluorescent GFP allele under the P*_flaB_* promoter ^21^. Strains were routinely cultured in gelatin-free BSK-II complete medium (supplemented with 50 µg/mL of gentamicin) in microaerophilic and static conditions at 33°C with 2% CO_2_. Strains were maintained as glycerol stocks at -80°C.

### Monoclonal antibodies (mAbs) and OspA immune sera

OspA specific mAbs (except LA-2) used in this study (**Table 1**) were expressed and purified by Dr. Lisa Cavacini (UMass Chan Medical School, Worcester, MA), as described ^15^. Chimeric LA-2 IgG1 was custom synthesized by ZabBio, Inc (San Diego, CA) ^7^. These mAbs were previously classified into three non-overlapping ‘bins’, based on their epitope targets on OspA ^6, 15, 16, 22^. Mouse OspA immune sera were kindly provided by Dr. Meredith Finn (Moderna, Inc.) ^17^.

### *B. burgdorferi* preparation for hanging drop motility assay

Frozen glycerol stocks of respective *B. burgdorferi* strains were thawed and cultured for ∼3 days, then diluted in fresh medium at a starting cell concentration of 1.5×10^6^ spirochetes/mL and grown till mid-exponential phase (2-5×10^7^ spirochetes/mL; ∼40-42 hours). Spirochetes were collected by centrifugation at 1800×g at room temperature. The cell pellets were washed once with phosphate buffered saline (PBS), pH 7.4, and then resuspended in gelatin-free BSK-II complete medium, at a cell concentration of 10^8^ spirochetes/mL. After resuspension, culture worth 2.5×10^6^ spirochetes from each strain were mixed (1:1 ratio v/v), following which, an equal volume of 4% porcine skin gelatin (Sigma-Aldrich, St. Louis, MO) solution was mixed in a 1:1 (v/v) ratio with the total volume of mixed culture. 50 µL of this final mixture was used for treatment with each mAb or serum. Spirochetes were treated with 10 µg/mL final concentration of mAbs or 1:50 final dilution of sera and incubated for either <5 min, 30 min, or 60 min at ∼28°C.

### Slide preparation

For microscopy, 1 µL of each sample (untreated control or antibody-treated) was taken on the center of a 30 mm × 22 mm and 0.16-0.19 mm thickness coverslip (VWR, Randor, PA). The coverslip was then inverted carefully on a glass depression slide (VWR), while maintaining the drop turgor and ensuring that the culture on the cover slip stays as a hanging drop and does not spread out. Finally, the coverslip was sealed on all sides using VALAP (Vaseline: lanolin: paraffin at 1:1:1 w/w/w).

### Microscopy and image processing

Imaging was conducted using a Nikon ECLIPSE Ti2-E microscope, along with the NIS Elements software version 6.10.01 (Nikon Instruments, Melville, NY). For capturing bacterial motility at a single cell level, a 40x fluorescence oil objective (Evident Scientific, Needham, MA) was used, with an average exposure of 60 ms and timelapse of 2 min with 2 second intervals. Images were collected in the GFP/FITC/Cy2 channel (Ex 466/40 nm, Em 525/50 nm) for strain GGW 1082 and DSRed/TRITC/Cy3 channel (Ex 554/23 nm, Em 609/54 nm) for strain GGW 1073 and saved as ND2 files. Images were collected from at least two or more different fields of view on each channel per treatment.

For image processing and quantifying cellular motility, the ND2 files were converted to TIFF 8-bit type files with the Fiji Bio-Formats plug-in ^23^. Images were preprocessed uniformly before segmentation (gamma correction: 0.7; mean filter: 4.0) to allow for optimal threshold of visualization and noise reduction, respectively. The localized thresholding algorithm of Phansalkar ^24^ was applied with default settings (radius: 15 and parameters 1 and 2: 0), followed by application of “white objects on black background” on all stacks. The same preprocessing and thresholding parameters were applied across conditions within each experiment. After these adjustments, the TrackMate plugin ^25^ was located, and metadata from each image file was automatically extracted into the program. The Mask detector function ^26^ was selected to simplify contours, and the threshold was manually verified by visual inspection to ensure no artifacts were introduced (e.g., incorrectly merged or split cells, debris identified as cells). For tracking each cell, the LAP tracker function was used with default settings (Linking maximum distance: 15.0 pixels; Gap-closing maximum distance: 15.0 pixels; Gap-closing maximum frame gap: 2). After the tracking algorithm had completed, we focused on the ‘Tracks’ from the display options and exported the Track files as .csv for further analyses. Each output track reported the ‘track duration’ (≤ 2 min) for each spirochete in the field of view, the ‘total distance traveled’ in that duration (µm), and ‘track mean speed’ (total distance traveled/track duration; µm/s).

### Statistical analysis and replication

Because track duration varied among cells, track mean speed was treated as the primary quantitative readout of motility, whereas total distance traveled during each accepted track was reported as a complementary measure of productive translocation. These motility parameters were collected from the tracks of all combined fields of view and plotted as violin plots, where each data point represents the parameter of a single spirochete in a particular treatment. Parameters from each independent replicate of the treatments were averaged and plotted as columns, and appropriate statistical analyses were performed, using GraphPad Prism v11.0.0, by comparing the strains GGW1073 (OspA^+^) and GGW1082 (OspA^-^) in a single treatment, as well as computing differences between treatments and with respect to incubation time.

### Microsphere immunoassay (MIA)

Polyclonal sera from sham- or OspA mRNA-LNP-vaccinated mice were collected ∼21 days post *B. burgdorferi*-tick challenge {Finn, 2026 #8338}. Seroconversion (as a measure of infection status) and OspA antibody levels were determined by microsphere immunoassay (MIA) using MagPlex-C microspheres (5 µg antigen/1×10^6^ microspheres) coupled to recombinant OspA ST1, OspC_A_ and DbpB {Finn, 2026 #8338; Willsey, 2026 #8336} (Freeman-Gallant et al., *manuscript in revision*).

### Antibody conjugation and OspA tracking on spirochete surface

We conjugated LA-2 with the Alexa Fluor™ 488 Antibody Labeling Kit (Life Technologies Corporation, Eugene, OR), following manufacturer’s instructions. Briefly, 1 mg/mL antibody solution in PBS was mixed with 1 M sodium bicarbonate buffer. The solution was then mixed with the reactive dye and incubated for 1 hour at room temperature, after which it was purified through a resin spin column. GGW1073 (OspA^+^, expressing mScarlet) spirochetes were treated with 10 µg/mL of the purified antibody and imaged in a hanging drop in the DSRed/TRITC/Cy3 channel (Ex 554/23 nm, Em 609/54 nm) for spirochete mScarlet expression and GFP/FITC/Cy2 channel (Ex 466/40 nm, Em 525/50 nm) for fluorescence conjugated-LA-2 bound to OspA on spirochete surface at <5 min and 30 min of antibody treatment. Images were captured and analyzed in a similar manner as described previously, without the spirochete tracking. Images from both channels were overlaid using Fiji to track the localization of OspA on spirochete surface along with LA-2 induced changes in spirochete morphology.

## Author contributions

AB was responsible for methodology, investigation, formal analysis, visualization, and manuscript writing (draft and final). RY was responsible for methodology, investigation, formal analysis, and writing of the draft manuscript. TMB provided microscopy resources, software, methodology, and supervision. NM was responsible for conceptualization, project administration, supervision, manuscript writing (draft and final) and funding acquisition.

## Acknowledgements and funding

We would like to extend a special thanks to Richard Cole and Danielle Hunt of the Wadsworth Center’s Advanced Light Microscopy core for assistance in imaging set-up. We thank the Wadsworth Center’s Media and Tissue Culture for BSK II media and buffers used in the study. In addition, we are grateful to Dr. Graham Willsey (Wadsworth Center) for providing bacterial strains and Dr. Meredith Finn (Moderna, Inc) for providing mouse OspA antisera. We also thank Grace Freeman-Gallant for running serological analyses with the OspA antisera. This work was supported by the National Institute of Allergy and Infectious Diseases (NIAID), National Institutes of Health, Department of Health and Human Services, Contract No. 75N93019C00040 (PI/PD Mantis). This content is solely the responsibility of the authors and does not necessarily represent the official views of the NIH.

## Video Captions

**Video 1:** Untreated OspA^+^ (GGW1073) and OspA^-^ (GGW1082) strains at <5, 30, and 60 min timepoints. In the PDF version of this article, please click anywhere on the figure or caption to play the video in a separate window.

**Video 2:** Pinwheel motion of a single OspA-(GGW1082) spirochete. In the PDF version of this article, please click anywhere on the figure or caption to play the video in a separate window.

**Video 3:** Looping motion of a single OspA^-^ (GGW1082) spirochete. In the PDF version of this article, please click anywhere on the figure or caption to play the video in a separate window.

**Video 4:** Saltatory motion of a single OspA^-^ (GGW1082) spirochete. In the PDF version of this article, please click anywhere on the figure or caption to play the video in a separate window.

**Video 5:** Reversal motion of a single OspA^-^ (GGW1082) spirochete. In the PDF version of this article, please click anywhere on the figure or caption to play the video in a separate window.

**Video 6:** Burst motion of a single OspA^+^ (GGW1073) spirochete. In the PDF version of this article, please click anywhere on the figure or caption to play the video in a separate window.

**Video 7:** Single cell masking and tracking of OspA^+^ (GGW1073) spirochetes in untreated and LA-2-treated conditions. In the PDF version of this article, please click anywhere on the figure or caption to play the video in a separate window.

**Video 8:** OspA^+^ (GGW1073) and OspA^-^ (GGW1082) strains treated with 221-7 mAb for <5, 30, and 60 min. In the PDF version of this article, please click anywhere on the figure or caption to play the video in a separate window.

**Video 9:** OspA^+^ (GGW1073) and OspA^-^ (GGW1082) strains treated with 857-2 mAb for <5, 30, and 60 min. In the PDF version of this article, please click anywhere on the figure or caption to play the video in a separate window.

**Video 10:** OspA^+^ (GGW1073) and OspA^-^ (GGW1082) strains treated with 221-20 mAb for <5, 30, and 60 min. In the PDF version of this article, please click anywhere on the figure or caption to play the video in a separate window.

**Video 11:** OspA^+^ (GGW1073) and OspA^-^ (GGW1082) strains treated with 212-55 mAb for <5, 30, and 60 min. In the PDF version of this article, please click anywhere on the figure or caption to play the video in a separate window.

**Video 12:** OspA^+^ (GGW1073) and OspA^-^ (GGW1082) strains treated with 319-28 mAb for <5, 30, and 60 min. In the PDF version of this article, please click anywhere on the figure or caption to play the video in a separate window.

**Video 13:** OspA^+^ (GGW1073) and OspA^-^ (GGW1082) strains treated with LA-2 mAb for <5, 30, and 60 min. In the PDF version of this article, please click anywhere on the figure or caption to play the video in a separate window.

**Video 14:** OspA^+^ (GGW1073) and OspA^-^ (GGW1082) strains treated with 319-44 mAb for <5, 30, and 60 min. In the PDF version of this article, please click anywhere on the figure or caption to play the video in a separate window.

**Video 15:** OspA^+^ (GGW1073) and OspA^-^ (GGW1082) strains treated with 3-24 mAb for <5, 30, and 60 min. In the PDF version of this article, please click anywhere on the figure or caption to play the video in a separate window.

**Video 16:** OspA^+^ (GGW1073) and OspA^-^ (GGW1082) strains treated with serum from sham vaccinated mice for <5 and 30 min. In the PDF version of this article, please click anywhere on the figure or caption to play the video in a separate window.

**Video 17:** OspA^+^ (GGW1073) and OspA^-^ (GGW1082) strains treated for <5 and 30 min with serum from mice vaccinated with 1 µg dose of OspA mRNA-LNP vaccine. In the PDF version of this article, please click anywhere on the figure or caption to play the video in a separate window.

**Video 18:** OspA^+^ (GGW1073) and OspA^-^ (GGW1082) strains treated for <5 and 30 min with serum from mice vaccinated with 0.1 µg dose of OspA mRNA-LNP vaccine. In the PDF version of this article, please click anywhere on the figure or caption to play the video in a separate window.

**Video 19:** OspA^+^ (GGW1073) and OspA^-^ (GGW1082) strains treated for <5 and 30 min with serum from mice vaccinated with 0.05 µg dose of OspA mRNA-LNP vaccine. In the PDF version of this article, please click anywhere on the figure or caption to play the video in a separate window.

**Video 20:** OspA^+^ (GGW1073) and OspA^-^ (GGW1082) strains treated for <5 and 30 min with serum from mice vaccinated with 0.01 µg dose of OspA mRNA-LNP vaccine. In the PDF version of this article, please click anywhere on the figure or caption to play the video in a separate window.

**Video 21:** Motility arrest of antibody-treated OspA^+^ (GGW1073) spirochete coincides with the formation of a lariat. In the PDF version of this article, please click anywhere on the figure or caption to play the video in a separate window.

**Video 22:** OspA tracking on the surface of GGW1073 (OspA^+^; mScarlet expression) with LA-2 mAb tagged with Alexa Fluor™ 488 (green fluorescence). In the PDF version of this article, please click anywhere on the figure or caption to play the video in a separate window.

